# A Systematic Analysis of Read-Across Adaptations in Testing Proposal Evaluations by the European Chemicals Agency

**DOI:** 10.1101/2024.08.29.610278

**Authors:** Hannah M. Roe, Han-Hsuan D. Tsai, Nicholas Ball, Fred A. Wright, Weihsueh A. Chiu, Ivan Rusyn

## Abstract

An important element of the European Union’s “Registration, Evaluation, Authorisation and Restriction of Chemicals” (REACH) regulation is the evaluation by the European Chemicals Agency (ECHA) of testing proposals submitted by the registrants to address data gaps in standard REACH information requirements. The registrants may propose adaptations, and ECHA evaluates the reasoning and issues a written decision. Read-across is a common adaptation type, yet it is widely assumed that ECHA often does not agree that the justifications are adequate to waive standard testing requirements. From 2008 to August 2023, a total of 2,630 Testing Proposals were submitted to ECHA; of these, 1,538 had published decisions that were systematically evaluated in this study. Each document was manually reviewed, and information extracted for further analyses. Read-across hypotheses were standardized into 17 assessment elements (AEs); each submission was classified as to the AEs relied upon by the registrants and by ECHA. Data was analyzed for patterns and associations. Testing Proposal Evaluations (TPEs) with adaptations comprised 23% (353) of the total; analogue (168) or group (136) read-across adaptations were most common. Of 304 read-across-containing TPEs, 49% were accepted; the odds of acceptance were significantly greater for group read-across submissions. The data was analyzed by Annex (i.e., tonnage), test guideline study, read-across hypothesis AEs, as well as target and source substance types and their structural similarity. While most ECHA decisions with both positive and negative decisions on whether the proposed read-across was adequate were context-specific, a number of significant associations were identified that influence the odds of acceptance. Overall, this analysis provides an unbiased overview of 15 years of experience with testing proposal-specific read-across adaptations by both registrants and ECHA. These data will inform future submissions as they identify most critical AEs to increase the odds of read-across acceptance.

## Introduction

Testing Proposals are a critical element of the European Union’s (EU) “Registration, Evaluation, Authorisation and Restriction of Chemicals” (REACH) regulation [Article 40(2)] (European Council, 2006) insofar they are a mechanism to eliminate unnecessary testing in animals and other model organisms and ensure that the most appropriate tests are performed. These submissions are prepared by the registrants where they identify data gaps in complying with the standard information requirements for their registration type (ECHA, 2011). They provide the European Chemicals Agency (ECHA) the opportunity to comment on the proposed studies and to suggest refinements, so the information obtained from any new testing is most informative for hazard and risk characterization. Annex XI of REACH provides a range of possible adaptations to the standard information requirements (European Council, 2006). Among potential adaptations, read-across is one of the major methods used to fulfil information requirements in REACH (ECHA, 2023a). The Organisation for Economic Co-operation and Development (OECD) and ECHA published guidance on read-across (ECHA, 2015, 2017; OECD, 2017) and there are many authoritative commentaries form diverse stakeholders on ways to improve read-across in regulatory submissions (Ball et al., 2016; Pestana et al., 2022; Patlewicz et al., 2014; Blackburn and Stuard, 2014; Rovida et al., 2020).

Few studies exist that systematically evaluated regulatory decisions and the reasoning that regulators used to accept or reject read-across hypotheses. A review of published decisions by ECHA that were available as of July 2015 was a product of multi-stakeholder collaboration (Ball et al., 2016). Both compliance checks (CCH, 524 documents) and testing proposal evaluations (TPE, 388 documents) were examined with regards to the relative successes and pitfalls of different scientific arguments used in proposed read-across hypotheses. The timing of that publication coincided with the publication of the first edition of ECHA’s read-across assessment framework (RAAF) (ECHA, 2015); additional guidance (ECHA, 2017, 2022, 2020) and recommendations (https://echa.europa.eu/recommendations-to-registrants) has been provided by ECHA. The study by Ball et al (2016) summarized the state of the art in read-across based on their analysis of the data available then, and highlighted the areas where improvements were needed in improving justifications and informing the registrants about best practices and successful cases. It was also acknowledged that RAAF guidance was likely to have a major impact on read-across adaptations and that additional systematic analyses will be needed.

Another example is a recent systematic analysis of read-across in REACH registration dossiers, data was extracted for target-source analogue pairs for mono-constituent substances from the information in IUCLID (International Uniform Chemical Information Database) (Patlewicz et al., 2024). The authors looked only at the data provided in the submission dossier and did not consider if ECHA has evaluated these through the process TPE or CCH. They also focused on the entries where read-across was used to satisfy information requirements for repeated dose toxicity or developmental toxicity studies – standard test requirements for high-tonnage substances that require the use of a large number of animals (Taylor et al., 2014b; Rovida et al., 2023). The analyses were restricted to substances with defined organic structures, the final dataset comprised 270 target substances and 259 source substances. The study focused on examining physicochemical, structural and metabolic similarity between source and target substances, as well as on the analysis of dose-response data from the animal studies – comparing the data from IUCLID to predictions using the United States Environmental Protection Agency (US EPA) generalized read-across (GenRA) approach (Helman et al., 2019). This study concluded that identification of suitable analogues for read-across is not only a challenge with respect to defining a similarity cutoff, but also with respect to finding the substances that already have data that satisfy standard test requirements. Collectively, the study found that low structural similarity was a common occurrence in REACH submissions based on read-across. They also concluded that GenRA provides more conservative estimates for dose-response analysis of chemical effects.

Overall, the discussions on the best practices to perform and evaluate read-across are critical to increase the familiarity with this approach, and for strengthening the justifications for read-across adaptations in regulatory submissions. Active discourse between the industry and regulators continues and progress is being made to increase regulatory acceptance of read-across submissions. Concomitantly, it is important to determine what do successful examples of accepted read-across adaptations look like and whether there are common themes in accepted and rejected submissions to ECHA. The extensive database of ECHA decisions on both CCH and TPE submissions now exists because nearly 15 years elapsed since the first implementation of review of submissions under REACH regulation. The current study focused on TPE decisions that were public as of August 2023 – a total of 2,630 Testing Proposals were submitted to ECHA from 2008 to 2023; of these, 1,538 had published decisions and were evaluated herein. Each document was manually reviewed, and information extracted in a systematic approach. Read-across hypotheses were standardized into 17 assessment elements (AEs), and each submission was classified based on these AEs. Data was analyzed for patterns, including Annex (i.e., tonnage), test guideline study, read-across hypothesis AEs, and the structural similarity of target and source substances.

## Methods

### Review of ECHA TPE Decisions and Data Extraction

ECHA publishes its decisions on the testing proposal evaluation (TPE) on its website (https://echa.europa.eu/information-on-chemicals/dossier-evaluation-status). On August 11, 2023, we exported information on the identities of the substances listed (i.e., the target substance), stage of their evaluation process, and the date the decision (if any) was issued. Among all downloaded records, 2,636 records were identified as TPEs, of which 1,538 included a link to a publicly available decision. This information is listed as part of **Supplemental Table 1**.

All TPEs with available decisions (n=1,538) were downloaded from the abovementioned website in a PDF (portable document format) and manually evaluated for information on the registration and missing data required to address data gaps for evaluation under REACH. It should be noted that while PDF files are machine-readable, about half of the files we examined were scanned images of varying quality. Attempts to perform optical character recognition processing of these files resulted in poor text quality that would make machine reading difficult to impossible. In addition, the terminology used by ECHA, as well as document formats, have changed over time. Collectively, attempts to automatically process these documents for information retrieval were vacated in favor of manual examination of each document. Information extracted from each PDF at this stage included Annex, the proposed tests to fulfill the data gap, and ECHA’s decision on the proposal (for more detail, see **Supplemental Table 1**). This step included identifying documents in which adaptations (e.g., read-across) were proposed; ECHA’s decision(s) on the proposed adaptation was also recorded. Of the 1,538 TPEs examined, 310 documents were identified as containing read-across adaptations. However, when evaluating testing proposal with read-across, some documents were found to include more than one read-across (either a hypothesis or end-point). In total, 314 records were included in the final analysis.

Next, the documents that included read-across adaptations were evaluated in greater detail and additional information was extracted. First, we recorded the type of read-across, analogue or group/category, as well as the EC number for each source substance(s). The EC numbers for the target and source substances were used to search the ECHA database to identify the type of each substance from the registration dossier. ECHA classifies substances into three primary types – mono-constituent, multi-constituent, or substances of “unknown or variable composition, complex reaction products, or biological” (UVCB). There were instances where the substance type could not be ascertained from the TPE because it was redacted, or the registrant had ceased manufacturing and the registration dossier was no longer publicly accessible. We also recorded separately what OECD test(s) were proposed by the registrants to be adapted through read-across and ECHA’s decision on each adaptation. Second, to enable evaluation of the read-across hypotheses in each TPE, we defined 17 assessment elements (AEs, **Table 1**) based on (i) information in ECHA’s RAAF guidance documents (ECHA, 2015, 2017) and (ii) additional considerations presented in TPEs that were not part of RAAF. The AEs were grouped into three categories; those based on toxicodynamic considerations, on toxicokinetic considerations, and other assessment considerations. Additional details on each AE, including corresponding explanation from RAAFs and/or TPE decisions (where applicable) can be found in **Supplemental Table 2**. For each TPE, AEs were recorded separately as those proposed by the registrant as part of their read-across hypothesis, and those used by ECHA in justifying their decision. Third, for each read-across decision, we noted whether submission was based on analogue or group/category, and whether the proposed standard test requirement adaptation was found by ECHA to be satisfactory or not.

**Table 1.** Assessment Elements (AE) used in Testing proposal Evaluation (TPE) submissions and decisions. See **Supplemental Table 2** for a detailed explanation of each AE.

| Assessment Elements Based on <u>Toxicodynamic</u> and Other Considerations |  | Assessment Elements Based on <u>Toxicokinetic</u> Considerations |  |
| --- | --- | --- | --- |
| AE.1 | Structure/Physicochemical Properties | AE.9 | Formation of Common (Identical) Compounds |
| AE.2 | Common Biological Targets with Common Metabolites | AE.10 | Exposure of Biological Targets to Common Compounds |
| AE.3 | Common Biological Targets without Common Metabolites (*Qualitative*) | AE.11 | Extent of the Bioavailability of the Parent Compound |
| AE.4 | Common Biological Targets without Common Metabolites (*Quantitative*) | AE.12 | Formation and Impact of Non-common Compounds/Exposure to Other Compounds than those Linked to the Prediction |
| AE.5 | Environmental Degradation to Non-common Compounds | AE.13 | Metabolites of Source and Target have been Identified |
| AE.6 | Environmental Bioaccumulation of Potential Non-common Compounds | AE.14 | Potential Presence of Uncharacterized Metabolites or Impurities |
| AE.7 | Common Environmental Degradation Pathways | **Other Assessment Elements** |  |
| AE.8 | Toxicodynamic Similarity based on the Data from a Bridging (OECD Test Guideline) Study | AE.15 | Characterization of Constituents (for Multi-Constituent or UVCB Substances) |
|  |  | AE.16 | Toxicodynamic Similarity based on the Data from a Bridging (Not a Guideline) Study |
|  |  | AE.17 | Lack of Observed Adverse Effects |

### Statistical Data Analysis

Statistical analyses were performed using *R* 4.40 and GraphPad Prism (v. 10.2.2, GraphPad Software, La Jola, CA). Using predictors such as OECD test and the proposed AEs in testing proposals and decisions, we constructed 2×2 tables (Acceptance/Rejection of a testing proposal vs. other binary predictors). These tables were analyzed using Fisher’s exact test (`fisher.test`) for *p*-values, odds ratios, and associated confidence intervals. As these predictors were considered to be of independent interest, nominal confidence intervals and *p*-values are reported. Multiple logistic regression using `glm()`function in R was performed using submission year, decision year, group vs. analogue-based read-across, and each AE. For several AEs, the small number of associated testing proposals could produce unstable estimates resulting in infinite standard errors, and these AEs were removed in a single iterative step. Reported *p*-values for the multiple logistic regression were corrected using the Benjamini-Hochberg *q*-value via `p.adjust` and *q*<0.05 was considered as significant.

### Analysis of Chemical Similarity

For testing proposals with analogue read-across for mono-constituent target and source substances, the EC number of each substance was used to obtain “simplified molecular input line entry system” (SMILES) identifiers for subsequent analysis of structural similarity. For each substance, SMILES were used to obtain Morgan Fingerprint information using the “*rcdk*” R package (Guha, 2007) to calculate the similarity between source and target substances. The Jaccard Similarity Score (Chung et al., 2019) was derived for each source and target compound pair. A t-test was then performed to compare the Jaccard similarity scores between accepted and rejected cases. To account for the possibility that acceptance rates might differ for scores greater than a threshold, a rigorous cutpoint analysis was performed to find an “optimal” threshold, while accounting for the implicit multiple testing in such an analysis. Here permutation-based significance testing was employed as follows. First, Jaccard similarity scores were screened to identify an optimal cutpoint. Each possible cutpoint (defined as an observed Jaccard similarity score in the dataset) was evaluated, and the cutpoint that returned the smallest nominal *p*-value using Fisher’s exact test (acceptance/rejection vs. Jaccard similarity < or ≥ cutpoint) was identified as optimal. Second, a null population was constructed by permuting the accepted and rejected labels 100,000 times, and for each permutation these labels were combined with the observed Jaccard Similarity Scores. Finally, an empirical *p*-value was determined by computing the proportion of permuted minimum nominal *p*-values that were ≤ the minimum nominal *p*-value from the first step, and the final odds ratio using the associated optimal cutpoint.

## Results

We first examined all publicly accessible TPE decisions by ECHA (1,538 were public as of August 11, 2023) with respect to whether any adaptations were proposed (**Figure 2**). We found that over ¾ of TPEs (1,185 or 77% of all TPEs) did not include adaptations, i.e., the registrants acknowledged that they need to perform additional tests as a remedy to addressing the data gap(s) in standard information requirements for their substance(s). The remaining 353 testing proposal submissions (23% of total) contained one or more adaptations, most of these (86% or 304) contained some form of read-across reasoning for one or more tests. Among the remaining 49 (14%) testing proposals, 39 contained adaptations other than read-across, such as QSAR or Weight of Evidence; these were not evaluated because they were not based on read-across. Another 10 submissions contained read-across adaptations but were not examined herein because ECHA rejected these submissions as redundant (e.g., required data was already available in another registration or endpoint proposed not necessary based on tonnage band) without weighing in on the registrant-proposed read-across arguments (for details, see **Supplemental Table 3).** Among 304 examined TPEs with read-across adaptations, more than half (55%) were of the analogue type – ECHA accepted read-across justifications for 39% of these submissions. Among testing proposals that proposed the group type of read-across, a larger fraction (62%) was found to be acceptable. Overall, based on the published TPE decisions from 2008 to 2023, the odds that a testing proposal with read-across hypothesis would be found adequate by ECHA for group read-across submissions were 2.6 times as large as that for analogue read-across submission (p<0.05, Fisher’s exact test). It should be noted, however, that this result should be interpreted with caution because if a read-across hypothesis/justification was found to be satisfactory for a group of similar substances, then several “positive” decisions might ensue thus amplifying the numbers as compared to analogue-type read across submissions.

**Figure 1.**
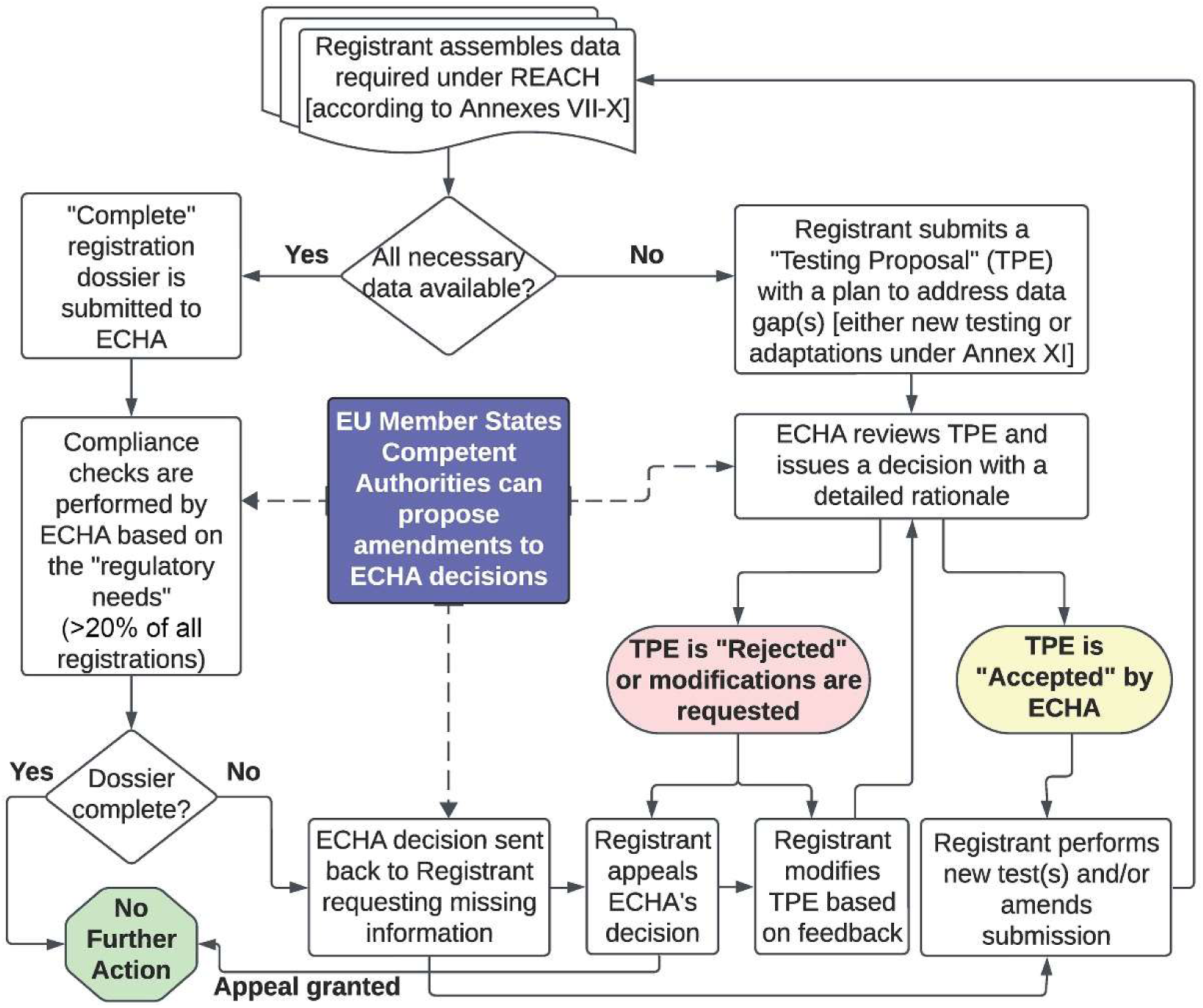
A simplified workflow of substance registration under REACH.

**Figure 2.**
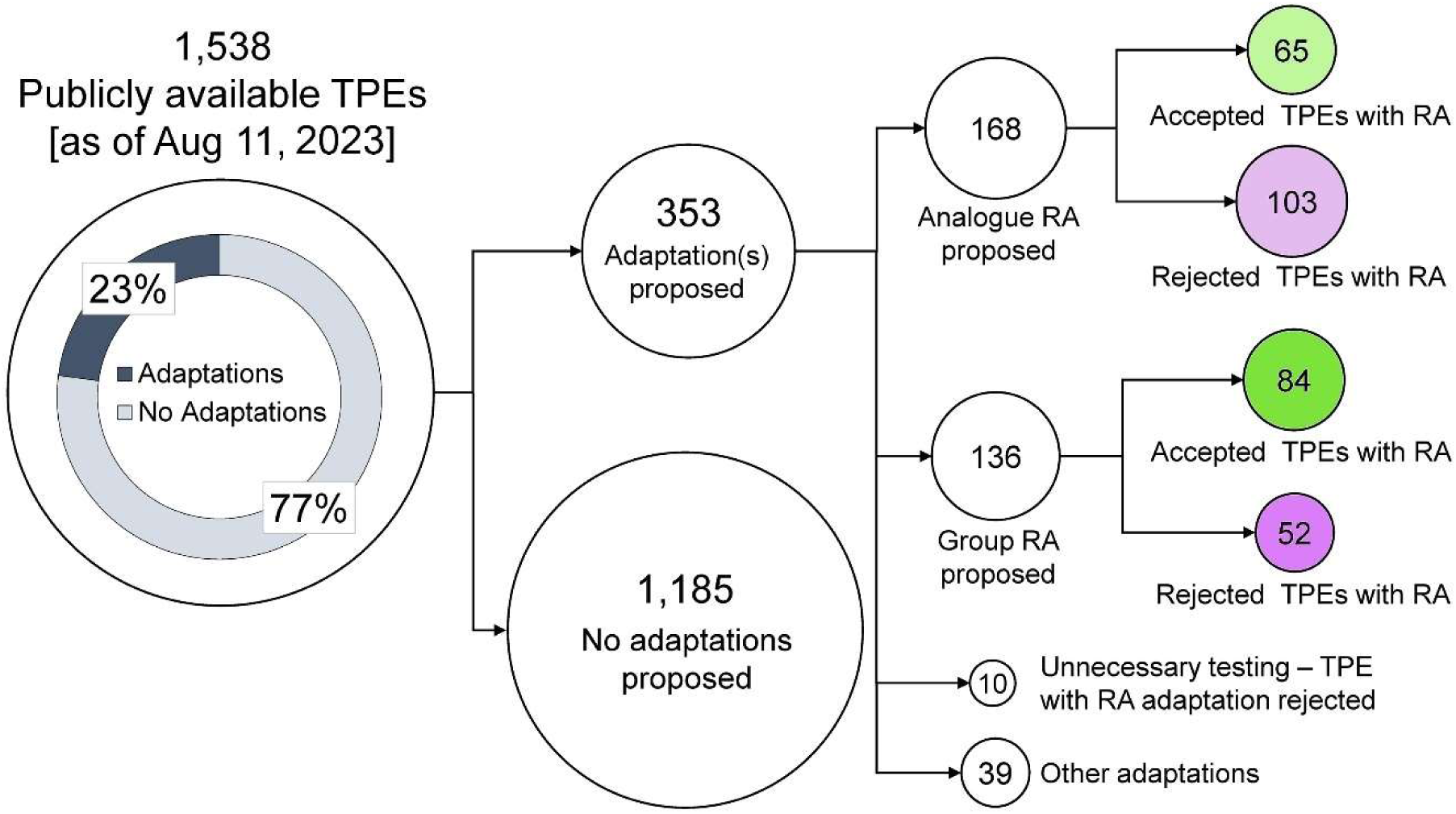
A diagram illustrating categorization of the publicly available TPE (Testing Proposal Examination) documents with and without adaptations, as well as the acceptance and rejection rates of TPEs with read-across (RA) adaptations. Numbers indicate the number of submissions with a published TPE as substances may have submitted multiple TPs throughout the registration period.

The process of evaluation of testing proposals by ECHA has been characterized as “lengthy and bureaucratic” (Taylor et al., 2014b), yet ECHA had to rapidly implement a substantial new regulatory framework, as well as establish internal expertise and competencies for evaluating a large number of testing proposals and registrations. **Figure 3A** shows the time trends in the public release of TPE decisions (bars) and the fraction of TPEs with read-across adaptations (line). Starting in 2012, when the output became more uniform, between 58 and 194 (126 ± 42, mean ± S.D.) TPE decisions were published by ECHA each calendar year. The fraction of TPEs with read-across adaptations varied even more widely, from 6.2% to 42% (22.5 ± 11.2%). **Figure 3B** shows the data for published TPEs with read-across adaptations. The stacked bars show the numbers of proposals with analogue and group read-across types. The line shows the fraction of read-across-containing submissions that were deemed acceptable by ECHA. There appears to be a large difference in the rate of acceptance with much higher acceptance rate in 2012-2015, before ECHA published final guidance on read across (ECHA, 2015). For example, in 2014, ECHA released decisions on a group of “Higher Olefins” substances which contained 21 substances. The decrease in acceptance of testing proposals between 2015 and 2017 may reflect the time needed to standardize the evaluation process according to RAAF. Overall, the time-dependent trends for the rate of testing proposal acceptance when either proposal submission year or decision publication year is considered, are slightly negative (slopes of -.078 and -0.101, respectively) but not significant.

**Figure 3.**
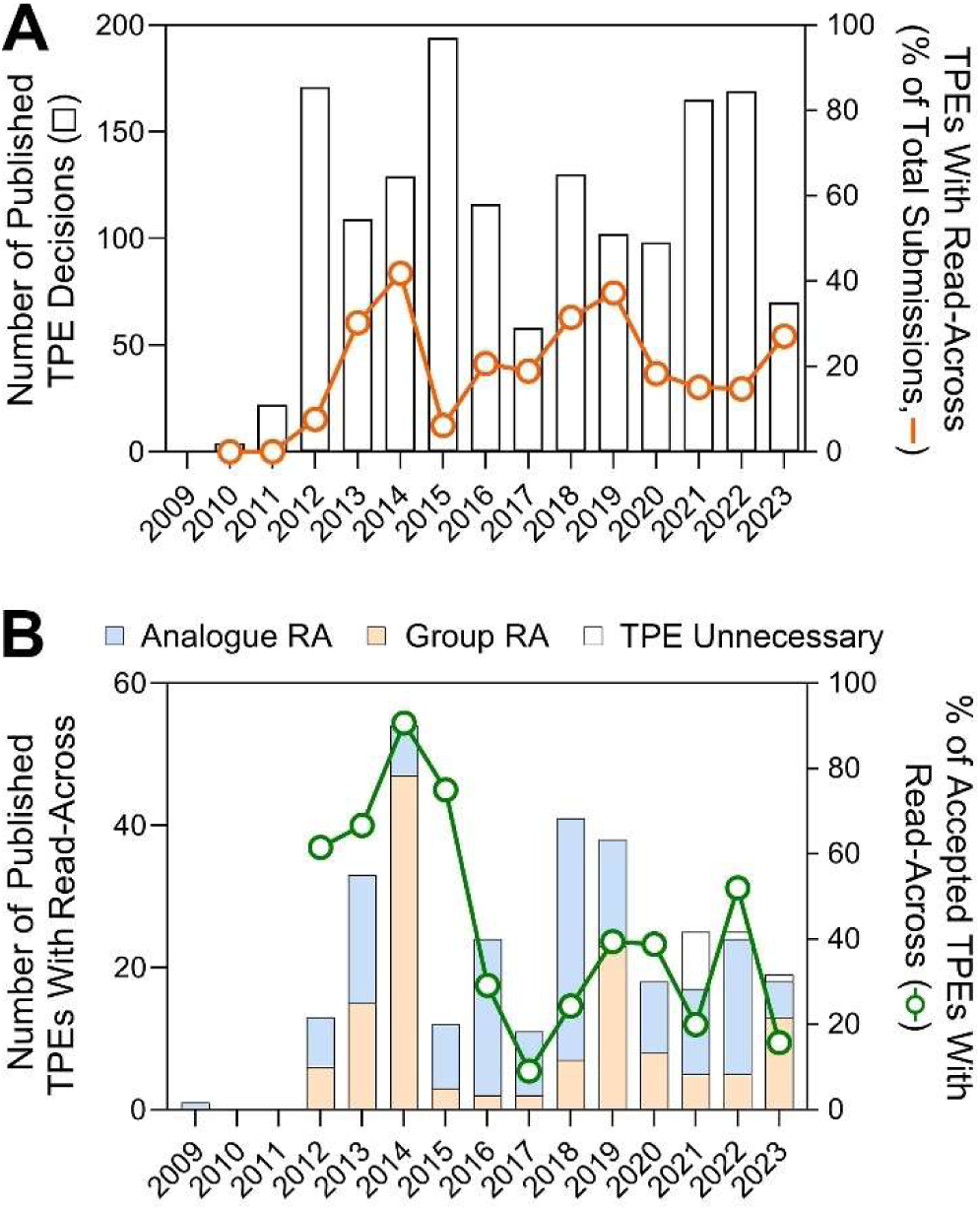
Time-trend plots for when TPE decisions were published. **(A)** The number of all published TPE decisions per calendar year (bars, left y-axis) and a fraction of TPEs that contained read-across (RA) adaptations (line, right y-axis). **(B)** The number of published TPE decisions per calendar year indicating the type of read-across (stacked bar plots, left y-axis) and a fraction of these that was accepted (line, right y-axis).

There are three main substance types recognized under REACH – mono-constituent, multi-constituent, and UVCBs [unknown or variable composition, complex reaction products or of biological materials] (ECHA, 2023b). **Figure 4A** shows that over 90% of published TPE decisions were on mono-constituent and UVCB substances. While the relative proportion of accepted read-across adaptations is about 50% for both, analogue submissions were most common for mono-constituent substances, while for UVCBs it was the group-type read-across that dominated. While the numbers for multi-constituent substances are relatively small, most submissions contained the analogue type read-across and the majority of these were not accepted. **Figure 4B** shows what source substance types were used by the registrants in read-across adaptations for each target substance type. While it is not surprising that most (79% for mono-constituent and 75% for UVCB) of the time the target and source substances were of the same type, it is curious that in some instances the registrants attempted to read-across from a UVCB to a mono-constituent substance. It also appears that when a UVCB substance was used as a source for another UVCB, more instances of the read-across adaptation were deemed inadequate as opposed to when a mono-constituent substance was used as the source; however, the differences were not significant between source/target pairs for either substance type.

**Figure 4.**
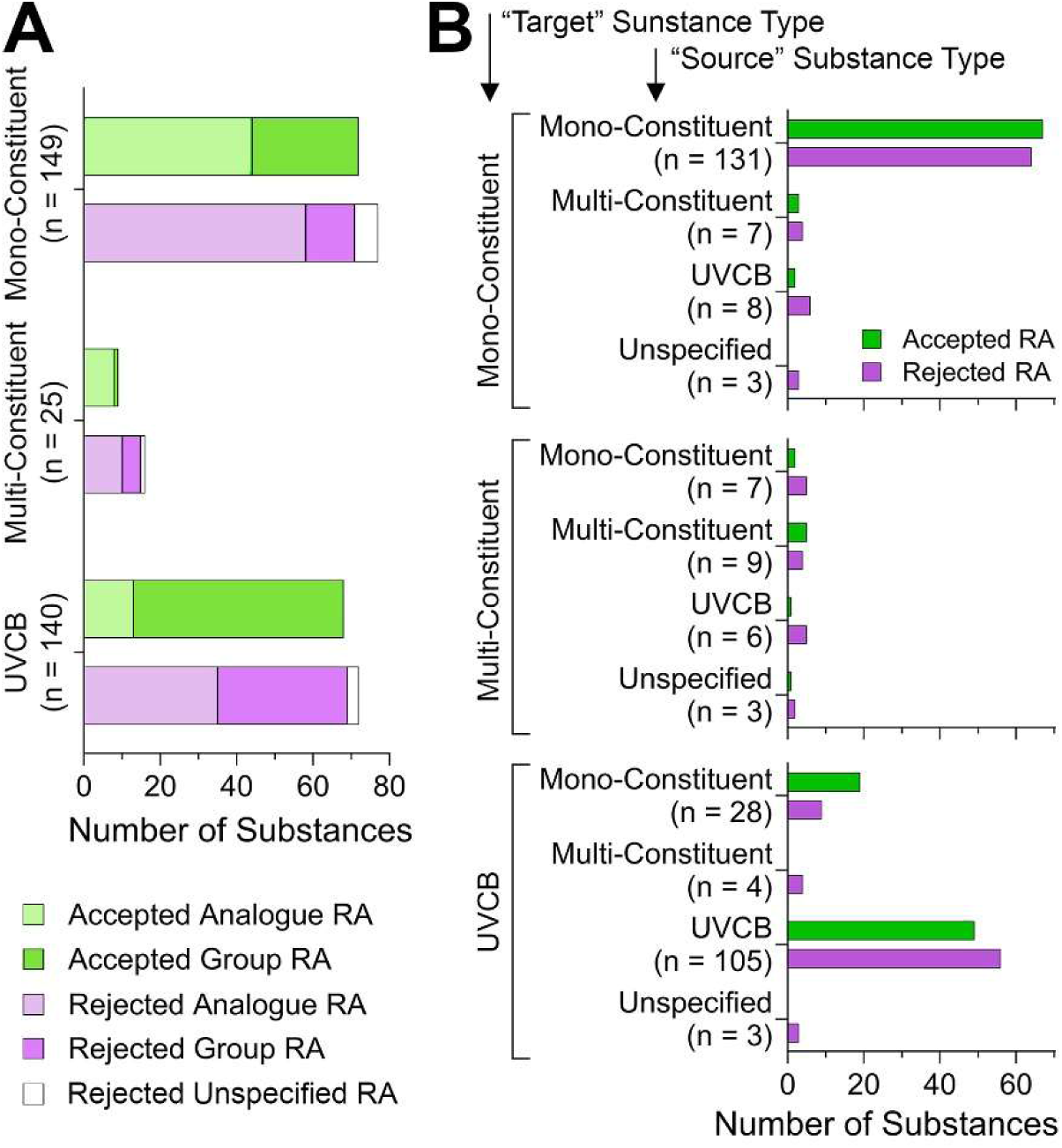
Analysis of the publicly available TPE decisions by substance type. **(A)** Stacked bar plots show the number of substances for which read-across adaptations were accepted or rejected, separated into substance categories. The total number of substances in each category is shown. Within each stacked bar plot, submissions that used analogue (lighter shade) or group (darker shade) read-across are shown. In some instances of rejected read-across, the type of read-across could not be determined (white). **(B)** For each target substance type, different types of source substances were used as indicated. The outcome of read-across evaluation is shown by the adjacent bars.

The REACH regulation established information requirements for substance registration (ECHA, 2011); these are based on the annual tonnage produced or imported into the European Union – the higher the tonnage, the greater number of studies that must be done (Botham et al., 2023). **Figure 5** shows the number of read-across adaptations across tonnage bands, from Annex VII (1-10 tons) to Annex X (over 1,000 tons). Recent registration data from ECHA (ECHA, 2023a) shows that among ∼12,500 registered substances, most (39%) are of the lowest tonnage/data requirements. Substances that are subject to most comprehensive testing are in Annexes IX and X, these comprise about 19% of the total for each Annex (2,346 and 2,335, respectively). When substances with testing proposals are considered (**Figure 5A**), the trends are largely reversed – few Annex VII (3.1%) and VIII (9.8%) substances had published TPE decisions, while the bulk of the evaluations were for Annex IX (52.5%) and Annex X (34.6%) substances. All animal tests required by Annexes IX and X but not yet available require a testing proposal to be submitted, while Annexes VII and VIII only do under certain circumstances (ECHA, 2011). Most TPEs with read-across adaptations were for Annex IX; however, the relative proportion of such submissions is similar across four tonnage bands. When TPEs with read-across adaptations are compared by tonnage band, read-across type, and ECHA decision (**Figures 5B-C**), it is evident that most rejected testing proposals with read-across adaptations for substances in Annexes VIII and IX were of the analogue type. The analysis of the odds that a proposed read-across type would favor a positive outcome shows that indeed, the odds of acceptance are significantly greater for group read-across adaptation for Annex VIII and IX substances (**Figure 5D**). While the odds were highest for Annex VIII substances, the 95% confidence interval was also wide, owing to a relatively smaller number of observations.

**Figure 5.**
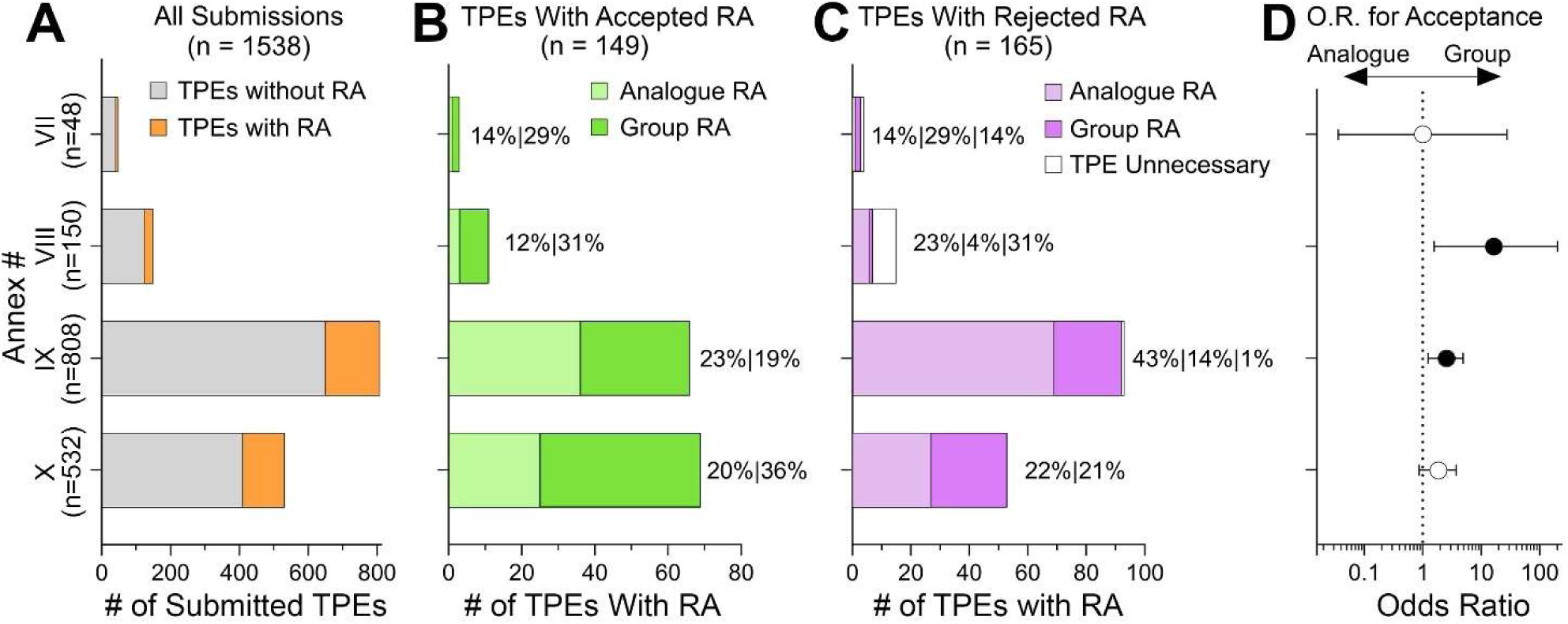
Analysis of the publicly available TPE decisions by REACH Annex number (substance tonnage). **(A)** Stacked bar plots show the number of substances in each Annex category for which publicly available TPEs were examined. Gray color represents TPEs without read-across adaptations and orange represents those with read-across adaptations. (**B-C**) The number of substances with accepted (B) and rejected (C) read-across adaptations. Within each stacked bar plot, submissions that used analogue (lighter shade) or group (darker shade) read-across are shown. In some instances of rejected read-across, the type of read-across could not be determined (white). **(D)** For each Annex number, odd ratios (OR) and 95% confidence intervals for Acceptance in Group vs. Analogue (i.e. an OR>1.0 corresponds to greater odds of Acceptance for Group). The intervals for Annex #IX and VIII do not contain the null OR=1.0, corresponding to a significantly greater odds of Acceptance for Group (*p*<0.05).

We also stratified the data by the type of a guideline test that was considered a data gap and where a read-across adaptation was proposed to fulfill data requirement(s) for registration under REACH. Most published TPE decisions concerned test of health effects in mammalian systems (**Figure 6A**). Among the so-called OECD test guideline (TG) 400 series assays which are designed to evaluate “health effects”, two assays predominated – TG 414 (Prenatal Developmental Toxicity Study) and TG 408 (Repeated Dose 90-Day Oral Toxicity Study in Rodents). **Figure 6B** shows the types of tests in OECD Guidelines for the Testing of Chemicals that are listed in sections 2 (Effects on Biotic Systems) and 3 (Environmental fate and behavior). Among these, the largest number of submissions was for TG 211 (Daphnia magna Reproduction Test). The fraction of submissions that involved read-across adaptations for tests with the greatest number of substances ranged from 14% to 29%. **Figures 6C-D** show the proportions of accepted and rejected TPEs for tests with read-across adaptations; most accepted read-across adaptations were of the group type. When the odds of acceptance for group vs analogue read-across were calculated, the significantly greater odds were for any test and TG 414 and TG 408 – a group read-across argument favors acceptance of an adaptation for these tests (for details, see **Supplemental Table 4**).

**Figure 6.**
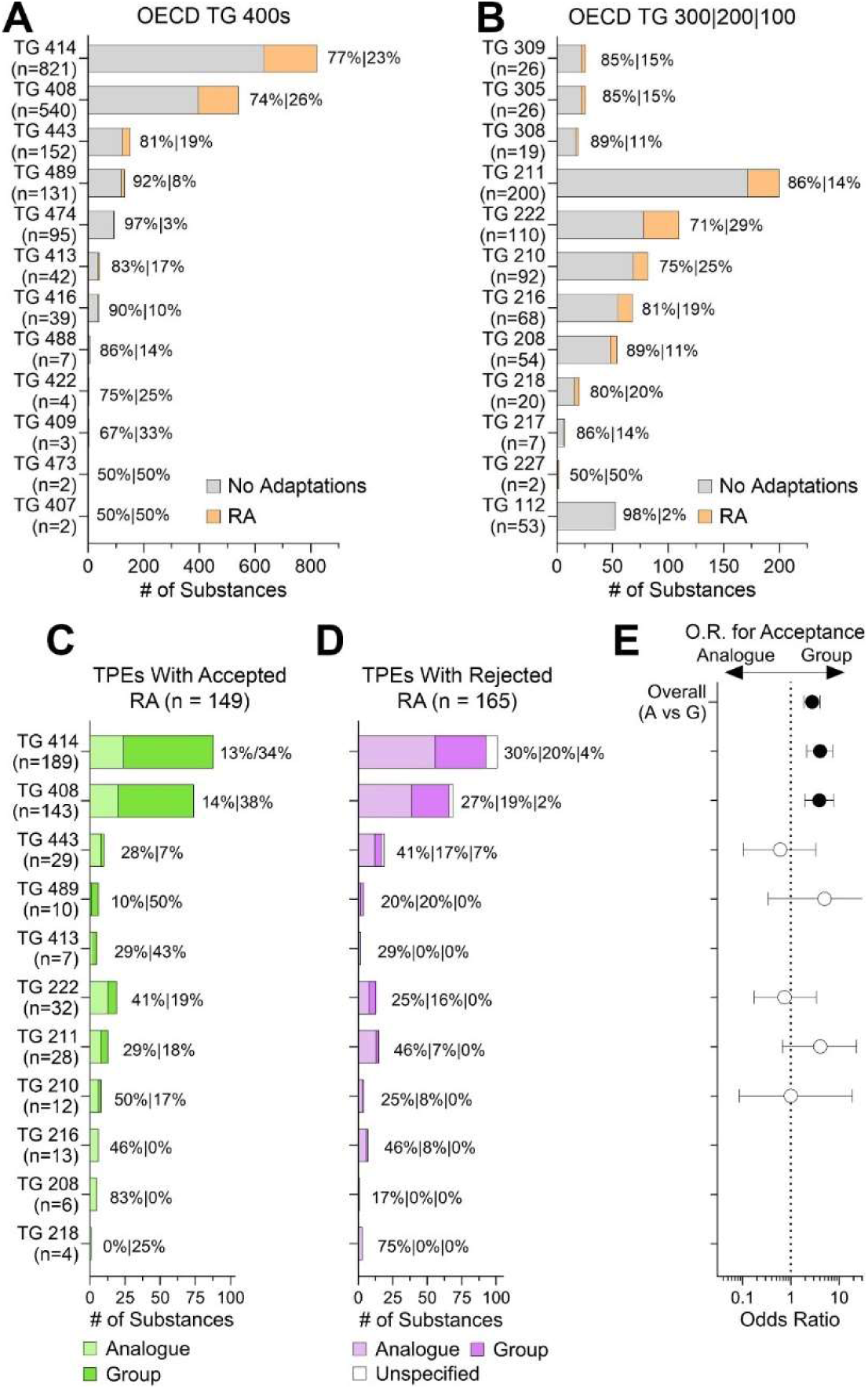
Analysis of the publicly available TPE decisions by OECD test guideline study type. **(A-B)** Stacked bar plots show the number of substances for each guideline test, separated into “human health” (A), and other (B) OECD test categories. Gray color represents TPEs without read-across adaptations and orange represents those with read-across adaptations. (**C-D**) The number of substances with accepted (C) and rejected (D) read-across adaptations. Within each stacked bar plot, submissions that used analogue (lighter shade) or group (darker shade) read-across are shown. In some instances of rejected read-across, the type of read-across could not be determined (white). **(E)** For the Overall data and when split by OECD guideline type, odd ratios (OR) and 95% confidence intervals for Acceptance in Group vs. Analogue (i.e. an OR>1.0 corresponds to greater odds of Acceptance for Group). The intervals for Overall, TG 414, TG 408, and TG443 do not contain the null OR=1.0, corresponding to a significantly greater odds of Acceptance for Group (*p*<0.05).

The type of read-across adaptation (analogue or group) may not be a matter of choice for each testing proposal depending on what source substance(s) with appropriate data are available. However, the registrants have options in building scientific support for their read-across hypothesis and can present a number of justifications based on ECHA guidance (ECHA, 2017, 2015, 2022, 2020) or other considerations (Ball et al., 2016; Beal et al., 2022; Berggren et al., 2015). It is generally accepted that a number of assessment elements, based on the considerations for how “similarity” is established between the target and source compounds, are needed to build the overall argument and justify the proposed adaptation. Because the decisions we evaluated spanned almost 15 years during which the best practices for read-across justifications have been evolving, we decided to craft “assessment elements” based on both formal ECHA guidance, as well as the arguments that were used in the decisions. The short list of assessment elements we used in the systematic analysis is shown in **Table 1**; additional information and example text from ECHA guidelines and decisions can be found in **Supplemental Table 2**. Each TPE decision document with read-across adaptation was examined with respect to what assessment elements were used by the registrants and separately by ECHA in their decision. Such analysis allows us to determine not only the frequency of use for each assessment element, but also the patterns of whether successful proposals have relied on a particular element or combination thereof.

**Figures 7A-B** show that among 17 possible assessment elements (AEs) we used in this evaluation, several have been used far more often than others. For example, arguments in support of structural/physicochemical similarity between target and source substances (AE 1) was included in virtually every TPE submission with read-across adaptation; however, there was no difference between successful and unsuccessful TPEs (for details, see **Supplemental Table 4**). Likewise, a large proportion of submissions relied on the reasoning that bridging studies from assays other than OECD test guidelines (AE 16) could aid in rationalization of the similarity argument; however, both successful and unsuccessful TPEs had them at the same rate. However, what is also evident from these figures is that the relative proportion of analogue vs group read-across was not identical between accepted and rejected TPEs. When odds of acceptance were analyzed for each assessment element with sufficient data (**Figure 7C**), we found that group-based read-across hypotheses that included AE 1, AE 8 (bringing studies that are OECD TG-type), and AE 15 (characterization of constituents for multi-constituent or UVCB substances) were significantly more likely to be accepted. **Figure 7D** shows the results from multiple logistic regressions, where submission year, decision year, group vs. analogue-based RA, and AEs were included in the same model. Overall, for a testing proposal with read-across to be successful, inclusion of AE 3 (qualitative identification of common biological targets without common metabolites) and AE15 significantly increased the odds. By contrast, inclusion of the information in support of AE 13 (identification of the metabolites of source and target) significantly increased the odds that the TPE will not be accepted. When the similar analyses were done separately for either mono-constituent substances, or UVCB and multi-constituent substances combined, the patterns were mostly similar but the significance for each assessment element was different. For mono-constituent substances, the overall direction was mostly the same as for all substances combined. However, the odds of rejection were considerable for UVCBs when AE 13 and AE 16 were part of a read-across hypothesis. Inclusion of the data showing quantitative evidence for common biological targets without common metabolites (AE 4) appeared to have a significant positive impact on the odds of acceptance, but this result is based on extremely few observations (only two within UVCB TPEs).

**Figure 7.**
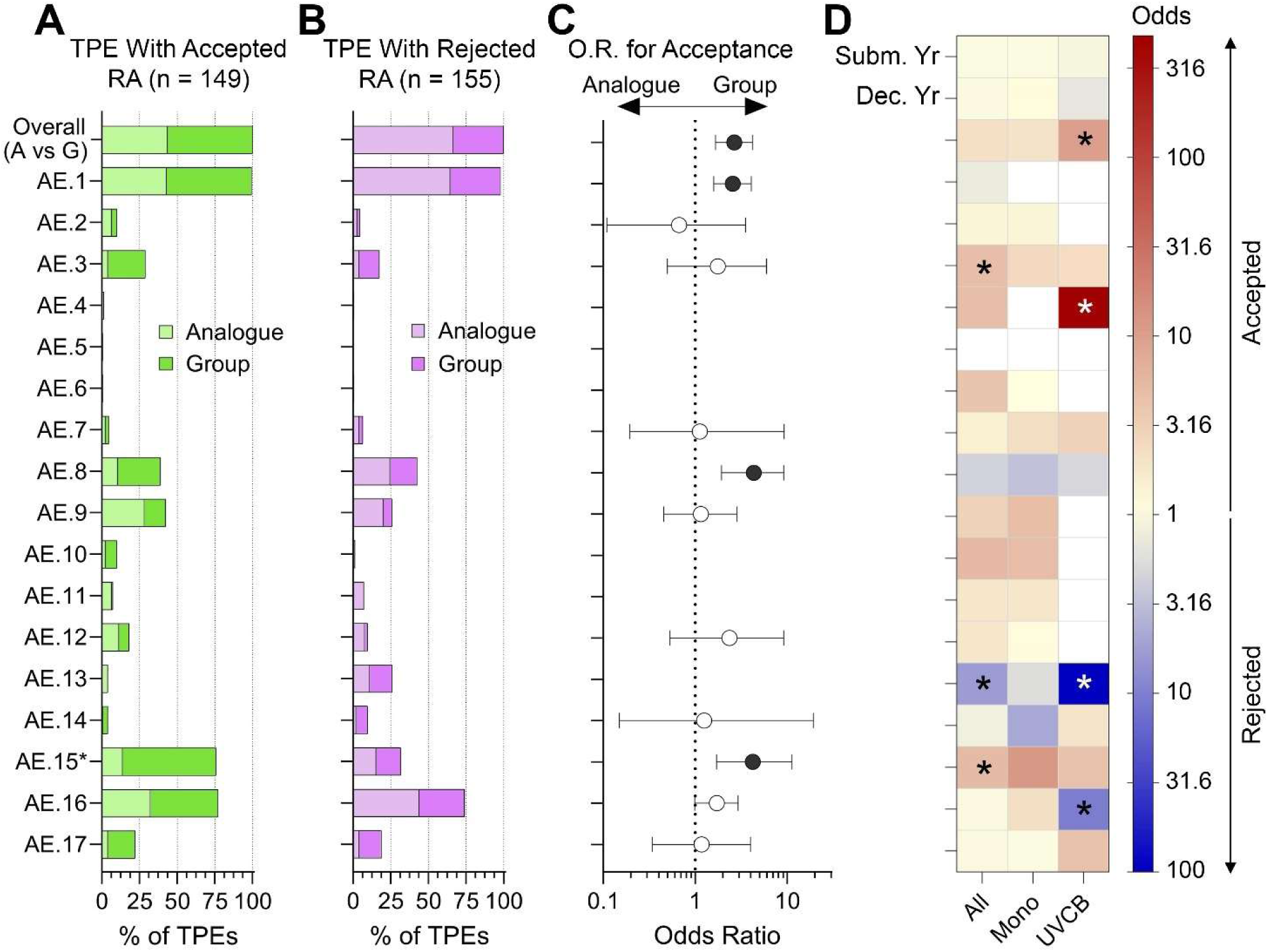
Analysis of the publicly available TPE decisions by assessment elements (AEs). **(A-B)** Stacked bar plots show the number of substances Overall and for each AE, separated into TPEs that were Accepted (A), and Rejected (B). For AE15 (marked with *), the fractions shown is for UVCB substances only. **(C)** For the Overall data and for each AE, odd ratios (OR) and 95% confidence intervals for Acceptance in Group vs. Analogue (i.e. an OR>1.0 corresponds to greater odds of Acceptance for Group). The intervals for Overall, AE 1, AE 8, and AE 15 do not contain the null OR=1.0, corresponding to a significantly greater odds of Acceptance for Group (*p*<0.05). **(D)** ORs from multiple logistic regression analyses with predictors submission year, decision year, group vs. analogue-based RA, and all AEs. For each column (all TPEs, Mono and UVCB), the ORs are displayed on a color scale, where red indicates predictor increases the chance of Acceptance, blue indicates predictor decreases chance of Acceptance, and ‘*’ denotes coefficients that have false discovery rates of *q*<0.05.

We also considered whether in unsuccessful TPEs with read-across adaptations (**Figure 7B**), the assessment element reasoning was presented by the registrants but not accepted, or if ECHA pointed out that certain types of data/reasoning were needed to accept the proposed adaptation. **Figure 8** shows the outcome of this analysis (for details, see **Supplemental Table 5**). In many instances, ECHA did not agree with the argumentation for AE 1 even though submissions contained information on this topic. The largest fraction of rejected submissions needed additional guideline bridging studies (AE 8), characterization of constituents (AE 15), or additional characterization of metabolites and/or impurities (AE 14). Characterization of the bioavailability of the parent molecule (AE 11) and quantitative analysis of common biological targets (AE 4) were frequently desired by ECHA but not provided by the registrants.

**Figure 8.**
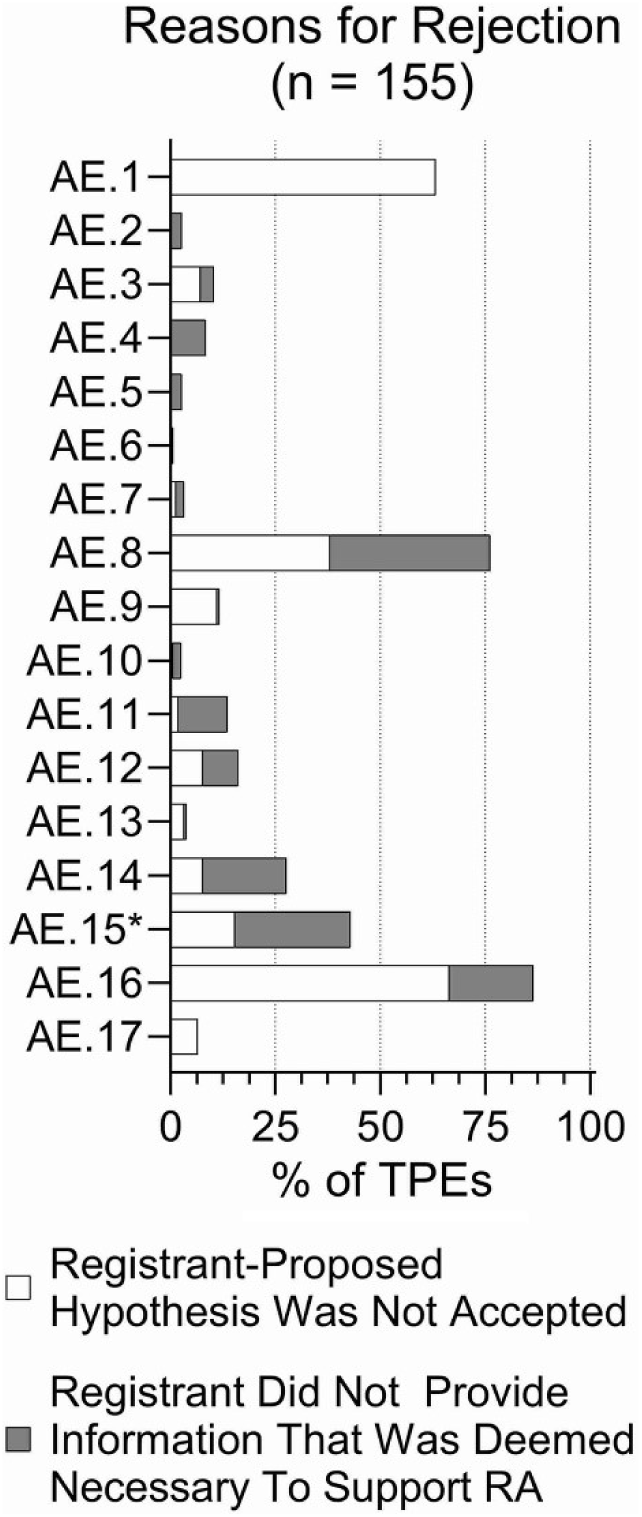
Reasons that were stated by ECHA for the rejection of a TPE with read-across. The light bar represents when a registrant proposed an element that ECHA disagreed with and/or interpreted differently. The dark bar represents the registrant failing to take an element into account in the read-across hypothesis.

Because AE 1 is the core element of building a similarity argument, we examined structural similarity between target and source compounds in TPEs with read-across adaptations. This analysis was restricted to mono-constituent substances for which chemical descriptors can be calculated, similar to the approach in (Patlewicz et al., 2024). In these analyses (**Figure 9**) we used structural similarity, based on Morgan fingerprints, and calculated Jaccard distance between target and source compounds as a metric for chemical similarity. Similar analyses were conducted using another type of chemical descriptors, the extensible chemistry-aware substructures called Saagar (Sedykh et al., 2021), and the results were nearly identical; therefore, we present here only the results of Morgan fingerprint analyses. For chemical pairs in TPEs with accepted read-across adaptations (**Figure 9A**), the greatest number were highly similar; however, there were many compounds that spanned the entire range. For example, the compounds with lowest similarity scores, but with accepted read-across (**Supplemental Table 6**), were metal-organic compounds such as Co^2+^ or Sr^2+^ 2-ethylhexanoate, Ba^2+^ or Co^2+^ carbonate, Pa^2+^ acetate ammoniate (1:2:4) and Co^2+^ acetate.

**Figure 9.**
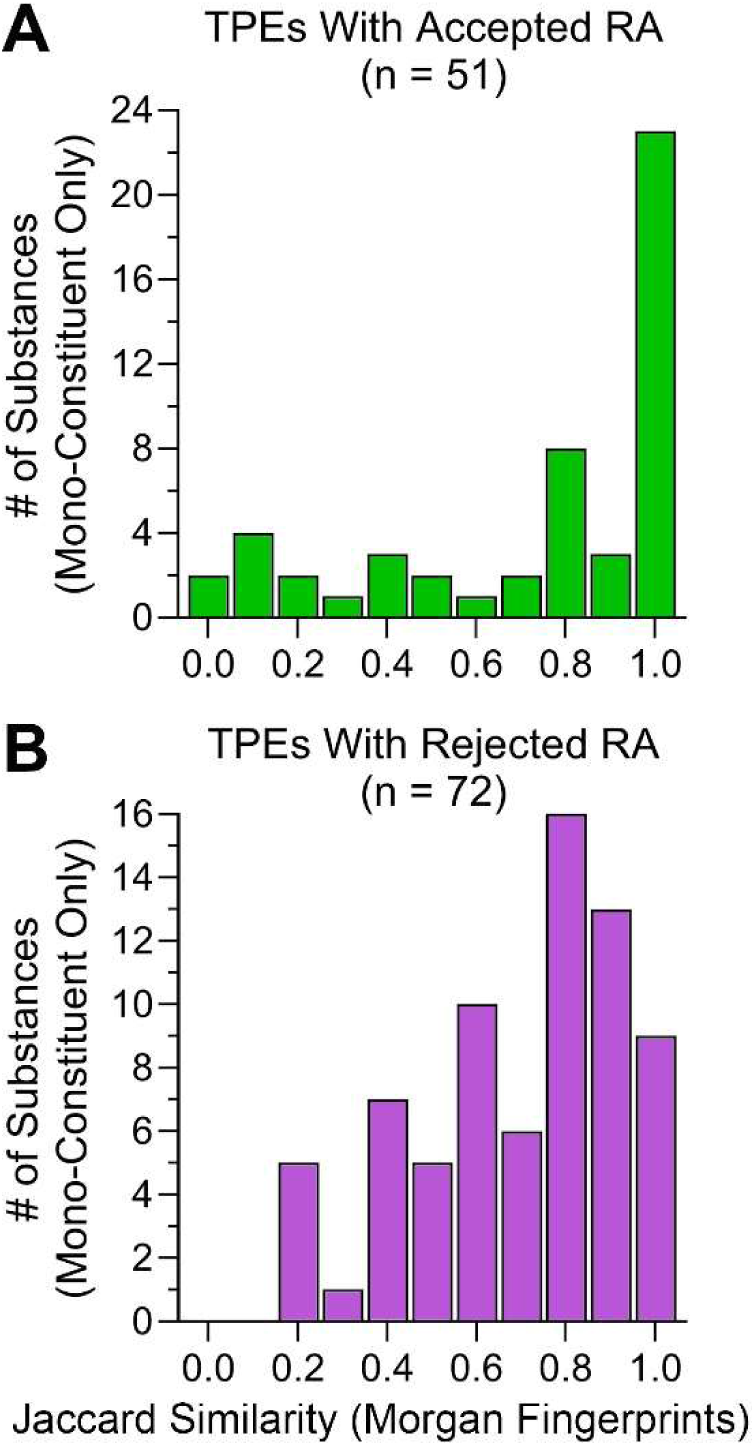
Source to target Jaccard similarity values for read-across proposals. **(A)** Accepted and **(B)** Rejected TPEs. Mean similarity values were not significantly different. However, a rigorous cutpoint analysis revealed that TPEs with very high similarity (>0.9) were significantly more likely to be accepted (p=0.0003, OR=7.0).

For substances in TPEs with rejected read-across adaptations, the distribution was much wider, but still right-skewed (**Figure 9B**). When the overall distributions in Jaccard similarity were compared between TPEs with accepted and rejected read-across adaptations, the difference in means was not significant (data not shown). However, when the analyses comparing Jaccard similarity between groups was restricted to substances with nearly perfect similarity based on Morgan fingerprints (score=1) and those with scores that were <1, then the chance of acceptance of the read-across adaptation was over 75% with a highly significant odds ratio of 7.0 (p=0.0003 via rigorous cutpoint analysis). When the similarity was not as high, the chance of acceptance was only 31%.

## Discussion

The goals of this study were three-fold – (i) to examine overall success rate of testing proposals that contained read-across adaptations to standard information requirements, (ii) to determine whether particular AE(s) made proposed read-across more or less successful, and (iii) to improve future read-across submissions by providing a structured database of information based on the actual ECHA decisions so that future proposals can identify relevant examples to “learn” from. As we mentioned in the Introduction, there have been many suggestions on how to “improve” read-across, often lamenting rigidities and conservatism of the decision-makers (Pestana et al., 2022; Patlewicz et al., 2014; Blackburn and Stuard, 2014; Rovida et al., 2020). However, relatively few studies (Ball et al., 2016) attempted to analyze the actual regulatory decisions on read-across submissions and identify specific areas where the registrants may need to do a better job in articulating their read-across justifications.

The study presented herein is a follow-up on the latter analysis, but almost a decade later and after numerous guidelines and recommendations were issued by ECHA in an attempt to interpret REACH and explain what justification are necessary for read-across hypotheses (ECHA, 2017, 2022, 2020, 2015). As our time-trend analysis showed, it does not appear that the success rates for read-across-based TPE submissions have been improving; therefore, the availability of guidance documents alone may not be sufficient to achieve success of successful adaptations. Similarly, it is also noteworthy that most for 77% of TPEs, the registrants did not propose adaptations and opted to just perform studies that were required. This can be either because there was no viable source compound(s), or because the registrants chose certainty with respect to REACH compliance, as opposed to the considerations of cost/time of doing such studies and/or animal welfare considerations. It is possible that the registrants were not interested in “trying” the adaptation route only to find that their submission was not acceptable, and that additional testing will still be necessary. Therefore, we reason that by examining the decisions and cataloging successful and failed read-across hypothesis it is possible to identify factors associated with positive decisions on read-across adaptations and to improve the outcomes of such submissions in the future.

### Overall Trends in Read-Across Adaptations in Testing Proposal Submissions and Decisions

There appears to be distinct trends in TPEs submissions with read-across – a rapid growth of read-across-containing adaptations as a fraction of all evaluated testing proposals before 2015, when the first RAAF guidance was published (ECHA, 2015), a precipitous decline to near zero in 2015, and another steep incline in the 5 years since 2015. On the decision side, a high proportion (60-90%) of read-across-containing TPEs submitted before 2014 was accepted; this was followed by a precipitous drop in 2016 and since. There are likely many reasons for these remarkable swings, most of these can be attributed to the registration deadlines, internal considerations at ECHA to deal with a large number of initial submissions under REACH, mutual learning of what acceptable read-across is, and intense advocacy from the industry to provide clear guidance on interpretation of vague language in REACH regulation. While certainly interesting from a historical perspective, and clear evidence that RAAF guidance did make an impact, perhaps not as intended, it is unlikely that a deeper dive into potential specifics of these trends will yield instructive learnings.

Our observation that the overall number of proposed adaptations was considerably greater for Annex IX and X substances is likely a direct result of increase in standard testing requirements with higher tonnage (Botham et al., 2023). Similarly, the finding that most common data gaps pertain to a few tests (e.g., TG 414 [Prenatal developmental toxicity study] and 408 [Repeated dose 90-day oral toxicity study in rodents]) is not surprising because these studies carry considerable cost and use a large number of animals (**Table 2**). For Annex X substances, the main human health information requirements are the TG 443 (Extended one generation reproduction study) and the TG 414 in a second species. Where a registrant required both, as well as the studies in Annex IX, it was common for several years for the TPEs to be split, with a decision on the TG 408 and first species in TG 414 and then follow up decision on the remaining studies once the results of the TG 408 and 414 were provided. This may explain why in general there were not as many TPEs covering these additional higher tier studies – because decisions on the follow up studies may still be under consideration and not publicly available decisions were available before this study’s cutoff of August 2023.

**Table 2.** Animal count for representative tests in animals that had proposed read-across adaptations in testing proposals to satisfy REACH substance registration requirements.

| OECD Test Guideline Studies That Were the Subject of Read-Across Adaptations in Testing Proposals | Accepted Testing Proposals with Read-Across: Number (% of 149 total) | Not Accepted Testing Proposals with Read-Across: Number (% of 165 total) | Number of Animals Required for Each Test |  |  |
| --- | --- | --- | --- | --- | --- |
|  |  |  | Minimum Number According to each OECD TG | Average Number According to (Taylor, 2018) | Average Number According to (Knight et al., 2023) |
| TG 408: Repeated Dose 90-Day Oral Toxicity Study in Rodents (OECD, 2018a) | 74 (49.7%) | 69 (41.8%) | 100 | 100 | 122 |
| TG 414: Prenatal Developmental Toxicity Study [rat] (OECD, 2018b) | 88 (59.1%) | 101 (61.2%) | 100* | 900 | 1,459 |
| TG 443: Extended One-Generation Reproductive Toxicity Study (OECD, 2018c) | 10 (7.1%) | 19 (11.5%) | 580 | 960 | 2,733/1,830<sup>#</sup> |
| TG 489: *In Vivo* Mammalian Alkaline Comet Assay (OECD, 2016) | 6 (4.0%) | 4 (2.4%) | 25 | - | 50 |

\*, Not including the number of animals in each litter.
<sup>#</sup>, Reflecting the use of mated/unmated animals.
-, Information was not included in the publication.

It is noteworthy that the apparent “success” rate of read-across adaptations was significantly different between analogue and group/category submissions. The odds for a successful testing proposal adaptation through group read-across were almost 3-times as great as those for the analogue read-across. These odds were most pronounced for the substances in Annexes VIII and IX, and for TG 414 and TG 408 studies. Even though this finding indicates that if the registrant had a choice between analogue and group, they may wish to opt into a group approach, there will be many instances where the choice of group or analogue read-across is dictated by the type of a substance and availability of the standard test data to which to read-across. In addition, one needs to keep in mind that for an analogue approach, one argument to use read-across covers one substance and so the ECHA decision covers just one substance. For the groups/categories, the same argument may cover multiple substances, but there will be a separate decision for each one, hence the appearance of a larger number of successful submissions may be misleading. Analogue-type submissions must prove that a source substance represents equal or worse case than target, while group-type submissions need to demonstrate a trend in the effect; therefore, it is generally more intuitive that read-across within a group/category would be more acceptable as compared to an analogue approach as there are more substances, and often more data from which to derive trends.

### Read-Across Assessment Elements – Examples

Similarity in structure/physicochemical properties (AE 1) is widely regarded as the foundational element for any read-across hypothesis and it is not surprising that this AE was part of virtually every read-across-containing testing proposal submission. For example, ECHA’s RAAF states that “*structural similarity is a pre-requisite for any grouping and read-across approach under REACH*” (ECHA, 2015). Similarly, this principle is central to the application of read-across to decisions by other agencies (Helman et al., 2019; Lizarraga et al., 2023); studies of the utility of various descriptors of chemical’s structure for prediction and characterization of chemical toxicity span several decades (McKinney et al., 2000). While structure-activity predictions are very useful, they also suffer from a number of pitfalls (Zvinavashe et al., 2008) and often need other data to increase confidence (Rusyn et al., 2012).

Our analysis of ECHA TPEs found that AE 1 was included in both accepted and rejected testing proposals, essentially to the same degree; however, very high chemical similarity (i.e., Jaccard similarity score of 1 based on Morgan fingerprints) was a strong predictor of acceptable read-across. Still, testing proposals with read-across for substances with very low structural similarity were also accepted. For example, there were several metalorganic compounds with acceptable read-across, but very low structural similarity based on Morgan fingerprints. One example is a group of “Cobalt-containing compounds” that included Co^2+^ 2-ethylhexanoate, Co^2+^ carbonate, and Co^2+^ diacetate. The registrants reasoned that a common metal cation, not the organic counterions, was the driver of any adverse health effects; the metal cation rapidly dissociates from the organic counterion when encountering the biological fluids. These submissions also included AE 9 (formation of common/identical compounds) and AE 12 (formation and impact of non-common compounds/exposure to other compounds than those linked to the prediction) as part of their read-across hypothesis. The overall rationale presented in these testing proposals was found acceptable by ECHA. It is interesting that decisions on other metalorganic compounds that used similar arguments were published at different times between 2013 to 2023, demonstrating that well-rationalized reasoning based on AE 9 and 12 can overcome low similarity in AE 1.

It should be noted that while considerations of toxicokinetics (AE 9 through 14) are widely regarded as important for “good” read-across (Ball et al., 2016; Rovida et al., 2020), there were relatively few submissions that included AEs other than 9 and 12, as discussed above. In fact, inclusion of AE 13 (Metabolites of source and target have been identified) to argue for exposure to structurally similar metabolites or different compounds that cause the same effect appeared to greatly diminish the odds of acceptance. AE 13 was present in 4% of accepted and 26% of rejected testing proposals with read-across. ECHA used AE 13 as a reason to reject proposed read-across in 4% of all unfavorable decisions – 3% were a difference of opinion on how the data was interpreted or whether the data are supportive of the read-across hypothesis, and 1% was the lack of discussion of this element by the registrant. For example, when computational predictors of metabolism were used without corresponding analytical evidence, ECHA did not find those arguments satisfactory. ECHA also frequently noted the lack of discussion of other metabolites that could be formed (i.e., AE 12) and whether those may be associated with adverse health effects.

The second most common element in proposed read-across hypotheses was AE 16 (Toxicodynamic similarity based on the data from a bridging (not a guideline) study). This AE includes any non-TG data submitted to support a read-across, such as *in vitro* methods, QSAR, and non-guideline animal studies. It was proposed in 77% of accepted and 74% of rejected testing proposals with read-across. It is even more noteworthy that ECHA used this AE to reject a read-across adaptation in 86% of all unfavorable decisions. Among these, ECHA disagreed with the strengths of the registrant’s reasoning based on the available data in 66% of decisions; in 20% of decisions, ECHA pointed out that such data would be needed to strengthen the read-across hypothesis. For example, the use of short-term studies to justify read-across adaptation of chronic or prenatal developmental toxicity studies or presenting the data for only the target or source substance, or for a different endpoint, was not deemed to be a satisfactory justification. Similarly, when target and source substances did not demonstrate similar effects in non-guideline studies, or when observations in non-animal (e.g., *in vitro* test) studies were not substantiated by data from *in vivo* studies, the read-across hypotheses were found to be not acceptable. For example, the data from *in vitro* mouse lymphoma assay, without support from *in vivo* studies, is not adequate as a justification for adaptations of higher-tier endpoints (e.g., TG 408).

A related read-across hypothesis element was AE 8 (Bridging [guideline] study), for which the registrant submitted data derived from a test conducted using an OECD TG protocol. AE 8 was part of 39% of accepted and 43% of rejected testing proposals with read-across; odds of acceptable read-across adaptation were 4.3 for group-type submissions that had this AE as part of their hypothesis. ECHA decisions discussed AE 8 in 76% of unfavorable decisions; half of these cases were a disagreement with the registrant as to whether such data were supportive of the read-across hypothesis, and another half were cases where ECHA pointed out that such data would be needed to justify the proposed adaptation. For example, when testing proposals for higher-tier endpoints (TG 408 and TG 414) included data from TG 422 (Combined repeated dose toxicity with the reproduction/developmental toxicity screening) in support of read-across and showing similar effects (e.g., same target organ, magnitude of effects, etc.) for both source and target substance, ECHA generally agreed with the registrant’s hypothesis. It is also worth noting that AE 8 was frequently a focus of “third party” comments on testing proposal submissions. Specifically, “third parties” reasoned that the findings of low toxicity in a 28-day oral study (TG 407) should be used as “bridging” evidence in support of read-across for adaptation of the TG 408 (90-day study) (Taylor and Andrew, 2017; Taylor et al., 2014a). ECHA considered these arguments in their decisions; however, the Agency stated in their decisions that it is the registrant’s (i.e., and not “third party”) responsibility to consider such reasoning and any other justifications when proposing adaptations.

Two AEs pertain to the argument that compounds may have common biological effects even if they are not structurally similar or do not form identical metabolites. Specifically, AE 3 (Common Biological Targets without Common Metabolites (Qualitative)) and AE 4 (Common Biological Targets without Common Metabolites (Quantitative)) address this type of a rationale in read-across. While AE 3 was included in a fairly large number of testing proposals with read-across, AE 4 was only found in 2 submissions. While it is impossible to extrapolate from only 2 submissions with AE 4, both of those were accepted by ECHA, it very well may indicate that quantitative arguments (i.e., the magnitude of the effect) are welcome. The latter conclusion is also supported by the fact that ECHA pointed out the lack of information pertaining to AE 4 in many unfavorable decisions on submitted testing proposals with read across (13%). Reasoning related to AE 3 was included in 29% of accepted and 17% of rejected testing proposals with read-across. Among these, ECHA disagreed with the strengths of the registrant’s reasoning based on the available data in 11% of decisions; in 5% of decisions, ECHA pointed out that such data would be needed to strengthen the read-across hypothesis. For example, submissions based on large groups (e.g., higher olefins and resin acids) relied on the arguments pertaining to AE 3 when reasoning that “*different compounds have the same effect*.” The inclusion of AE 3 significantly increased the odds of acceptance for group-type submissions.

Substances that are classified as multi-constituent or UVCB have several additional challenges with respect to the need for establishing both substance identity, and to characterize the chemical composition to support read-across. ECHA published separate guidance for these substances (ECHA, 2017); in addition, the chasm between the regulator’s expectations and the realities of analytical chemistry solutions for identifying and quantifying the constituents in highly complex substances has been documented (Roman-Hubers et al., 2023). It is not surprising, therefore, that AE 15 (Characterization of multi-/UVCB substances) was commonly included in testing proposals with read-across adaptations for these types of substances. In submissions concerning multi-constituent/UVCB substances, AE 15 was present in 76% of accepted and 32% of rejected testing proposals with read-across. ECHA used this AE in 43% of unfavorable decisions; in 15% it was a difference of opinion on the interpretation of the available data and 28% because the registrant did not address this AE. The odds of acceptance were significantly higher (4.2) for group-type submissions when this AE was part of the read-across hypothesis. In the cases of unfavorable decision, ECHA reasoned that insufficient characterization was provided by the registrant for target or source (or both) substances, which means that there was no way to compare their similarity or lack thereof. While this reasoning is similar to that of AE 1, it is far less clear how to define “*broad similarity*” for substances that have too many constituents and when their composition is expected to be variable. Case examples of petroleum substances have been recently published with respect to the use of other supporting data types from *in vitro* studies to justify grouping (Tsai et al., 2023; House et al., 2022; House et al., 2021); however, ECHA did not find these non-guideline “bridging” studies (AE 16) satisfactory for a number of reasons. Many of these related to chemical characterization of the substances and/or their extracts that were used for *in vitro* testing, challenges that need to be addressed by additional research (Roman-Hubers et al., 2022; Cordova et al., 2022).

### Using a Database of TPE Read-Across Adaptations to Construct Future Submissions

Even though our analysis showed that about 50% of testing proposal submissions that relied on read-across adaptations were found to be acceptable by ECHA, improvements are needed to increase success. We found that over the past decade, the fraction of accepted proposals stayed relatively constant and the efforts by all parties involved in the registration process have yet to produce a measurable improvement. Therefore, we reason that the data we extracted from the decisions may provide instructive examples of successful submissions that could be emulated in the near future. It is also clear that by improving success rate of testing proposal submissions with read-across, measurable impact can be made in terms of reduction in animal use for chemical registrations under REACH. **Table 2** shows that the number of animals that will be required to meet information requirements if new testing is performed is substantial. Similarly, if more registrants will consider read-across-based or other adaptations, rather than defaulting to new testing, the reduction in animal use will be even more pronounced.

While it is without a doubt that the registrants and ECHA are committed to sharing encouraging examples and mutually develop best practices for read-across, we reason that the data collected in this project will prove useful to both registrants and regulators (for full database, see **Supplemental Tables 1, and 3-5**). On the one hand, to achieve greater chance of success, registrants will be able to identify specific aspects of read-across justifications that merit most attention and improve these in the testing proposals they are working on. Our results also likely to encourage consideration of a category versus analogue approach, although the latter may not be possible for many submissions. We also found that read-across between different substance types should be discouraged, particularly reading-across from UVCBs to mono-constituent substances. On the other hand, the trends identified in our study could indicate changes in ECHA’s approach to interpretation of read-across and suggest that additional efforts may be needed by ECHA to improve consistency in the application of their guidance over time. In addition, ECHA and other agencies may use our results and the database to identify areas where they could provide more granular advice to improve how read-across arguments are presented and justified.

### Limitations of the Study

While our analyses and interpretation of the findings may prove instructive, we also acknowledge that our results may not be taken as definitive and deterministic. First, we note that our dataset may have limitations as it had a cutoff date of August 2023 and the trends in most recent submissions, and thus decisions, may be different given the attention given to the efforts to improve read-across (Pestana et al., 2022; Patlewicz et al., 2014; Blackburn and Stuard, 2014; Rovida et al., 2020). Second, we relied on the robust summaries of the testing proposals as included in ECHA’s decisions, we did not examine the testing proposal submissions. While it is possible that some mis-interpretation could have occurred, we reason that robust summaries may be eventually more informative in terms of categorizing the elements of the proposed read-across into standardized categories (i.e., AEs, **Table 1**). Even though the AEs were defined by the authors, we reason that no single guidance, or combination of different documents released by ECHA over time, clearly defines read-across AEs. While this is another limitation, we point out that every effort was made to assure that the AEs used herein were comprehensive and consistent with both RAAF and the actual decisions (**Supplemental Table 2**).

Third, a large difference in the success rate of read-across adaptations between testing proposals and compliance checks (Ball et al., 2016) indicates that the “bar” may be lower on the former because the decision to allow for the use of read-across as part of a testing proposal is provisional. Whether the use of read-across is ultimately acceptable or not for the registration can only be decided once the registration dossiers are submitted. Consequently, there may be cases where the use of read-across has been accepted for the testing proposal but was subsequently challenged or rejected. For example, the registrants of decan-4-olide (EC: 211-892-8) submitted a testing proposal with read-across adaptation for TG 211 (long-term toxicity to aquatic invertebrates) and this submission was found to be acceptable by ECHA in 2015. However, a compliance check decision by ECHA (published in 2023) concluded a read-across adaptation to standard information requirement was insufficient and that required TG 211 data were needed to justify proposed classification. Even though this caveat does not invalidate the results presented in this manuscript, it does highlight that even “*final*” decisions on read-across can be subject to change.

Finally, we need to point out that although our analysis has identified some specific areas that were influential in read-across acceptance, it is rare that ECHA decisions focus on just one aspect; often, multiple reasons may lead to the ultimate decision to accept or not accept. Furthermore, the submission may be accepted not because ECHA agreed with the rationale presented in a testing proposal, but because different combination of the information included in the submission was deemed to be sufficient. For example, for the Asphalt UVCB (EC: 232-490-4) submission, the registrant proposed a read-across adaptation of TG 414 to another substance in the group (Residues (petroleum), thermal cracked vacuum; CAS: 92062-05-0) on which this test is to be performed. The principal reasoning by the registrant was that the target and source substances belong to a group formed based on the refining process and on similarity in carbon number distribution and the hydrocarbon class profiles. Furthermore, the registrant hypothesized that one of the hydrocarbon classes – polycyclic aromatic hydrocarbons containing 4 or more aromatic rings – is the putative reproductive toxicant. While ECHA found that the proposed justification for the overall group was insufficient, it agreed that a read-across to the proposed source substance selected based on the PAH with 4+ aromatic rings was “appropriate.” While such cases complicate the overall analysis as they make it difficult to be confident that one or two specific AEs had an impact on each decision, our data still identified several “critical” AEs as detailed above. Ultimately, we encourage the registrants to focus their efforts on improving the rationales for their read-across hypotheses to stand a better chance of acceptance. Even though each read-across submission is unique in terms of the type of a substance, availability of the data and the endpoint, we show that there are several general trends and that the registrants can rely on case examples in the database when crafting their submissions.

## Supporting information

All records from ECHA Website.xlsx

Details on Assessment Elements.xlsx

All Decisions

Proposed Assessment Elements.xlsx

Reasons for Rejection

Structural Similarity

## Data availability

Data extracted from publicly available testing proposal evaluations by ECHA are included as Supplemental Tables.

## Acknowledgements

The authors are grateful to ECHA Hazard Assessment Directorate staff for general advice and encouragement on this project. The authors would also like to thank Drs. Lauren Zeise, Kannan Krishnan, and Anatoly Soshilov at OEHHA for useful discussions.

